# Effects of *Pseudomonas aeruginosa* pyocyanin and 1-hydroxyphenazine on intracellular calcium, mitochondrial function, and viability in human nasal epithelial cells

**DOI:** 10.1101/2024.11.08.622701

**Authors:** Joel C. Thompson, April Park, Yobouet Ines Kouakou, Zoey A. Miller, Nabil F. Darwich, Nithin D. Adappa, James N. Palmer, Robert J. Lee

## Abstract

*Pseudomonas aeruginosa* is an opportunistic pathogen that produces phenazine metabolites pyocyanin and 1-hydroxyphenazine that have been suggested to have detrimental effects on mitochondrial function and reactive oxygen species (ROS) production. Prior studies have suggested activation of Ca^2+^ signaling by pyocyanin in an airway cell line, while others have shown apoptotic effects on cancer cells. Ca^2+^ is tightly linked to both normal mitochondrial function as well as mitochondrial ROS and apoptosis during mitochondrial Ca^2+^ overload. We found that pyocyanin but not 1-hydroxyphenazine induced both cytosolic and mitochondrial Ca^2+^ increases in RPMI 2650 and primary human nasal epithelial cells (HNEC). Similar results were seen in HNEC air-liquid interface (ALI) cells, but these cells did not display a cytosolic Ca^2+^ response after treatment with pyocyanin. In RPMI 2650, activation of PKC, ER Ca^2+^ release, and PLC inhibition, indicated potential GPCR activation from pyocyanin. Similarly, ROS production increased after treatment with pyocyanin, but not 1-hydroxyphenazine in all 3 cell types, but with stark differences in CF and non-CF ALIs. In HNEC ALI, pyocyanin reduced ciliary beat frequency (CBF) after 4 hours, while 1-hydroxpyhenazine did not. Despite this, both pyocyanin and 1-hydroxyphenazine decreased in cell viability in RPMI 2650 nasal carcinoma cells but not in HNEC at 24 hours. However, in both RPMI 2650 and HNEC, mitochondrial membrane potential acutely decreased after treatment with either pyocyanin or 1-hydroxyphenazine. Finally, 24-hour pyocyanin treatment decreased expression of ER stress genes in some cancer cells, but not in non-cancerous HNEC. Our data suggest that Ca^2+^ signaling is not required for acute effects of 1-hydroxyphenazine or pyocyanin on mitochondrial function. The greater sensitivity of RPMI 2650 cells to pyocyanin-induced and 1-hydroxyphenzine-induced cytotoxicity compared with primary cells suggests that these compounds might have some applicability in treating nasal squamous carcinomas or other types of head and neck squamous carcinomas. Although the exact mechanisms of pyocyanin induced apoptosis remains uncertain, the downregulation of the ER stress response may play a role.

## Introduction

*Pseudomonas aeruginosa* is an opportunistic pathogen which primarily infects immunocompromised individuals such as burn victims or cancer patients, and causes fatal respiratory infections in patients with cystic fibrosis^**1,2**^. The prevalence of multidrug-resistant *Pseudomonas aeruginosa* has increased in recent years, making it a significant public health threat in these patient populations^**3**^. Understanding the interactions of *P. aeruginosa* with host cells may aid the development of new therapies that might circumvent antibiotic resistance^**4**^. We previously studied activation of bitter taste receptors (taste family 2 receptors or T2Rs) by *P. aeruginosa* acyl-homoserine lactone (AHLs) and quinolone molecules involved in quorum sensing^**5**^. *P. aeruginosa* AHL, 3-oxo-C12HSL induces apoptosis in epithelial cells through activation of T2R14^**6**^, which causes mitochondrial Ca^2+^ overload and excessive mitochondrial reactive oxygen species production that inhibits the ubiquitin proteasome pathway^**7**^.

*P. aeruginosa* synthesizes multiple other toxic virulence factors^**8**^, including phenazine metabolites pyocyanin and 1-hydroxyphenazine, before and during biofilm development^**1,9**^. Concentrations of phenazines in late-stationary cultures or biofilms can reach between 100-300 µM^**10**^, and 100 µM pyocyanin has been reported in CF patient sputum^**11**^ and infected human ear secretions^**12**^. Pyocyanin is a “redox-active” compound that has been reported to induce cellular apoptosis through superoxide production^**1,2,13**^, and although the exact mechanisms are unclear, they likely involve mitochondrial dysfunction. One study showed that in human monocyte-derived macrophages, pyocyanin did not induce apoptosis via caspase 3/7 activation after 24 hours^**14**^. Other studies have shown that pyocyanin increases in activating transcription factor 6 (ATF6) in non-cancerous NRK-52E cells^**15**^, which plays a role in endoplasmic reticulum unfolded protein response (UPR^ER^) and could play a role in downstream endoplasmic reticulum-associated degradation (ERAD), although this has not yet been show. Pyocyanin has been suggested to have potential anti-cancerous effects via activation of apoptosis, and also has potential anti-fungal and anti-bacterial properties^**1,2**^.

We sought to better understand the role of Ca^2+^ in the response to pyocyanin and 1-hydroxyphenazine. Calcium is a master regulator of many cellular processes and has a biphasic relationship with mitochondria. While calcium is required for proper mitochondrial function, too much Ca^2+^ leads to cell death^**16,17**^. In A549 lung cancer cells, pyocyanin induced a cytosolic Ca^2+^ response in the presence and absence of extracellular Ca^2+^, suggesting endoplasmic reticulum (ER) Ca^2+^ release^**18**^, which would fit with activation of a G protein-coupled receptor (GPCR). Other studies have shown that pyocyanin and 1-hydrpoxyphenazine can diminish or even completely inhibit ciliary beating in human nasal epithelial cells^**19,20,21**^, as we have also observed with activation of T2R1 in nasal epithelial cells^**16**^. It is unknown if pyocyanin activates Ca^2+^ responses in primary airway epithelial cells, and if it does, if this is GPCR related. One goal here was to test if excessive mitochondrial Ca^2+^ influx is linked to mitochondrial dysfunction and epithelial damage, as we observed with another *P. aeruginosa* metabolite, 3-oxo-C12HSL^**6**^. Since only one study has investigated the effects of pyocyanin, albeit in a cancer cell line, and no studies have examined 1-hydroxyphenazine in Ca^2+^ signaling, these compounds warrant more investigation in primary airway epithelial cells. Defining the effects of these molecules on airway epithelial cells will lead to insight into Ca^2+^ signaling in the context of in host-pathogen interactions. Our goal was to further study host-pathogen interactions in human nasal epithelial cells mediated by pyocyanin and 1-hydroxyphenazine both because of their roles in infection as well as their potential utility as apoptosis-inducing cancer therapeutics. We also sought to determine if phenazines activated bitter taste receptors, as observed with AHLs, to understand if these molecules shared similar mechanisms of apoptosis induction. Because we also previously showed that T2R bitter taste receptor responses can be altered in primary nasal epithelial cells from people with CF^**22**^, a secondary goal here was to determine if cells from CF individuals had altered responses to pyocyanin or 1-hydroxyphenazine.

## Materials and Methods

### Cell culture

RPMI 2650 (ATCC CCL-30) and NCI-H520 (ATCC HTB-182) cells were from ATCC (Manassas, VA, USA), while UM-SCC-47 hereafter referred to as SCC-47 (Millipore SCC071) was from MilliporeSigma (St. Louis, MO, USA) and all were cultured as described^**23**^. Cells were cultured in Minimal Essential Medium with Earle’s salts and L-glutamine (Corning; Glendale, AZ, USA) with 10% (v/v) fetal bovine serum (FBS) (MedSupply Partners; Atlanta, GA, USA), 1% (v/v) penicillin/streptomycin (PenStrep) mix (Gibco; Gaithersburg, MD, USA), and 1% (v/v) MEM nonessential amino acids (Gibco). Cell media was changed twice per week and cells were passaged between 70-80% confluence.

Human nasal epithelial cells (HNEC) were extracted via cytology brushings which occurred during sinus surgery, as previously described^**24**^. Brushes were obtained from patients undergoing sinonasal surgery for CRS or other procedures (e.g., trans-nasal approaches to the skull base for tumor resection) with institutional review board (IRB) approval (protocol #800614) and with written informed consent. All work was done in accordance with The University of Pennsylvania guidelines for use of residual clinical material, the U.S. Department of Health and Human Services code of federal regulation Title 45 CFR 46.116, and the Declaration of Helsinki. Inclusion criteria were patients ≥18 years of age undergoing sinonasal surgery. Exclusion criteria included use of antibiotics, oral corticosteroids, or anti-biologics (e.g. Xolair) within one month of surgery or history of systemic inheritable disease, not including cystic fibrosis (e.g., granulomatosis with polyangiitis, systemic immunodeficiencies). Individuals ≤18 years of age, pregnant women, and cognitively impaired persons were excluded.

HNECs were isolated from cytology brushings by washing 5 times with DMEM/F12 media containing 5% (v/v) FBS, 1% (v/v) PenStrep, and 0.125 µg/mL amphotericin B (Gibco) referred to as Culture Media (CM). Cells were centrifuged at 1200 rpm for 5 mins, then combined in 5 mL CM and strained through a 70 µm mesh strainer (Fisher). Cells were centrifuged again using the same settings, and were resuspended in PneumaCult Ex-Plus Medium (StemCell Technologies; Vancouver, Canada) containing 1% (v/v) PenStrep, 0.125 µg/mL amphotericin B, 10 µM ROCK inhibitor Y-27632 (Tocris; Bristol, UK), and 1 µM DMH-1 (Tocris) referred to as Growth Media (GM). HNECs in the expansion phase (hereafter referred to as submerged HNECs) were cultured in T-25 flasks, had their media changed 2-3 times per week, and were passaged between 70-80% confluence. To get HNECs to differentiate, cells were seeded on Thincert (Grenier Bio-One, Kremsmünster, Austria), or Transwell (Corning) cell culture inserts (0.33 cm^**2**^ PTFE membrane, 0.4 μm pore size, transparent) and were grown, submerged in GM until cells were fully confluent. Once at confluence, cells were no longer submerged on the apical side, and were switched to PneumaCult ALI Basal Medium (StemCell Technologies) containing 1% (v/v) PenStrep and 0.125 µg/mL amphotericin B, referred to as Differentiation Media (DM) on the baso-lateral side only. Media was changed for these cells (hereafter referred to as HNEC ALI) 2-3 times per week and were allowed to differentiate for a period of at least 4 weeks, after first exposure to DM. Genotypes for CF patients used

### Reagents and Solutions

Hanks’ balanced salt solution (HBSS) was used in cell imaging experiments which contained 137 mM NaCl, 5 mM KCl, 400 µM KH2PO4, 300 µM Na2HPO4, with or without 5.6 mM glucose, 1.8 mM CaCl2, 1.5 mM MgCl2, 20 mM HEPES, pH 7.4. HBSS containing 5.6 mM glucose is hereafter referred to as HBSS with glucose, and HBSS without 5.6 mM glucose referred to as HBSS without glucose. Crystal Violet (CV) solution was used in cell viability assays which contained 0.1% CV in DI water with 10% acetic acid. Pyocyanin and 1-hydroxyphenazine were from Cayman Chemical.

### Live cell imaging of cytosolic Ca^2+^ (Ca^2+^_cyt_), mitochondrial Ca^2+^ (Ca^2+^_mito_), ER Ca^2+^ (Ca^2+^_ER_), and PKC activity

Submerged HNEC or RPMI 2650 cells were loaded with 5 µM Fluo-4 AM (AAT-bioquest; Pleasanton, California, USA) in HBSS with glucose for 1 hour in the dark at room temperature (RT). HNEC ALIs were loaded apically with 10 µM Fluo-4 AM in HBSS without glucose, containing 295 µg/mL Probenecid and 394 µg/mL Pluronic^®^ F127 for 2 hours at RT in the dark. For submerged cells (RPMI 2650 or submerged HNEC), cytosolic Ca^2+^ was imaged using a Nikon Eclipse TS100 microscope (20x 0.75 NA objective), FITC filters (Chroma Technologies), QImaging Retiga R1 camera (Teledyne; Tucson, Arizona, USA), MicroManager software, and XCite 120 LED Boost (Excelitas Technologies; Mississauga, Canada). For HNEC ALIs, cytosolic Ca^2+^ was imaged using an Olympus IX-83 microscope (10x 0.40 NA objective), FITC filters (Chroma Technologies), Orca Flash 4.0 sCMOS camera (Hamamatsu, Tokyo, Japan) and MetaFluor (Molecular Devices; Sunnyvale, CA USA)

For Ca^2+^_mito_, Ca^2+^_ER_, or PKC activity, submerged HNECs or RPMI 2650s were transfected with pCMV-Mito-LAR-Geco1.2 (Addgene Plasmid #61245;^**25**^), pcDNA-D1ER (Addgene Plasmid #36325;^**26**^), or CMV-ExRaiCKAR (Addgene Plasmid #118409;^**27**^) respectively, using Lipofectamine 3000 reagents (ThermoFisher Scientific) per the manufacturer’s instructions 48 hours prior to imaging. Ca^2+^_mito_ was imaged with an Olympus IX-83 microscope (20x 0.75 NA objective), TRITC filters (Chroma Technologies), Orca Flash 4.0 sCMOS camera (Hamamatsu, Tokyo, Japan) and MetaFluor (Molecular Devices; Sunnyvale, CA USA). Ca^2+^_ER_ or PKC activity was imaged as described above with the Olympus IX-83 microscope system, but with CFP and YFP, or TRITC and FITC filters (Chroma Technologies) respectively. For Ca^2+^_mito_ in HNEC ALI, cells were loaded with 5 µM Rhod-2 AM (AAT-bioquest;^**24,28**^) for 18 hours in the dark at 4°C. After loading, ALIs were incubated on the basolateral side only in F-12K Media (Gibco) at 37°C for 1 hour. Cells were imaged in HBSS without glucose on the apical side and HBSS with glucose on the basolateral side. Imaging occurred on the same system as HNEC ALI Ca^2+^_cyt_ but with a standard TRITC filter set and MetaMorph imaging software (Molecular Devices).

### Mitochondrial membrane potential, superoxide production, cell viability, and apoptosis measurement

For acute single-wavelength mitochondrial membrane potential measurement, HNEC (submerged or ALI) and RPMI 2650 cells were loaded for 15 minutes at RT in the dark with tetramethylrhodamine ethyl ester (TMRE) dye and Hoechst dye prepared as described in the Multi-Parameter Apoptosis Assay Kit (Cayman Chemical; Ann Arbor, MI, USA). Cells were imaged with TRITC filters (Chroma Technologies) as described above with the Olympus IX-83 microscope system and 20x lens (for submerged HNEC and RPMI 2650) or 10x lens (for ALI). For longer-term mitochondrial membrane potential measurement, submerged HNEC and RPMI 2650 cells were loaded for 15 minutes at 37°C and 5% CO_2_ with 1.5 μM JC-1 dye (Cayman Chemical), which allowed ratiometric green-red imaging. Imaging occurred as described above with the Olympus IX-83 microscope system, but with TRITC and FITC filters (Chroma Technologies) and MetaMorph (Molecular Devices).

For monitoring mitochondrial superoxide production, HNEC (submerged or ALI) and RPMI 2650 cells were loaded for 30 minutes at 37°C and 5% CO_2_ with 0.5 μM MitoSOX™ Red dye (Life Technologies; Carlsbad, CA, USA). Imaging occurred as described above with JC-1 imaging, but with TRITC filters (Chroma Technologies). For crystal violet cell viability assays, submerged HNEC and RPMI 2650 were treated, in their respective culture medias, for 24-hours. Cells remaining on plates were stained with the CV solution described above, then washed with DI water, and allowed to dry for 6-24 hours at RT. Stains were dissolved in 30% acetic acid in DI water and absorbance was measured in a Spark 10M Multimode Microplate Reader (Tecan; Männedorf, Switzerland) at 590 nm.

### Ciliary Beat Frequency (CBF)

To measure ciliary beat frequency, HNEC ALIs were equilibrated on the temperature control plate set at 37°C for at least 15 minutes. Equilibration occurred in HBSS without glucose on the apical side, and HBSS with glucose in the basolateral side. CBF was measured using Sisson-Ammons Video Analysis (SAVA) software^**29**^, a Nikon Diaphot microscope (Modulation Optics Inc. HMC 20x LWD 0.4NA 160/0-2 objective), and Basler acA1300-200um Camera (Ahrensburg, Germany) with a Nikon 0.7x DXM Lens (Tokyo, Japan), and Heating Insert M24 2000 EC (Pecon; Erbach, Germany). ALI inserts were treated apically, and CBF was recorded for 2.5 seconds each in 5-10 random locations per insert (averaged) every hour.

### Reverse transcription quantitative Polymerase Chain Reaction (RT-qPCR) expression of endoplasmic reticulum (ER) genes

RPMI 2650, SCC-47, NCI-H520, or HNEC ALI were treated, for either 16 hours, with various concentrations of pyocyanin 100 μM or 50 μM versus control (media). HNEC ALI samples were treated baso-laterally only. After exposure, cells from either type were lysed with TRIzol^®^ reagent (ThermoFisher Scientific) and RNA extraction was carried out using the DirectZol™ RNA Kit (Thermo Fisher Scientific; Waltham, MA) following the manufacturer’s instructions. Complementary DNA (cDNA) was synthesized using the High-Capacity cDNA Reverse Transcriptase Kit (ThermoFisher Scientific) in a Thermo-Cycler (Techne TC-512, Keison Products). RNA expression was quantified using TaqMan qPCR probes (QuantStudio 5 Real-Time PCR System, ThermoFisher Scientific). All expression levels were quantified using the 2^-ΔΔCT method and were compared to either UBC (RPMI 2650) or Rpl13a (SCC-47, NCI-H520, or HNEC ALI).

### Statistical Analysis

All data were analyzed for statistical significance using unpaired, two-tailed t-tests (2 comparisons) or one-way ANOVA (more than 2 comparisons) in GraphPad PRISM (San Diego, CA, USA). Dunnett’s and Bonferroni post tests were used for one-way ANOVA when appropriate as indicated in figure legends. In each of the figures, p < 0.05 (*), p < 0.01 (**), p < 0.001 (***), and no statistical significance (N.S or unmarked). All data points represent the mean ± SEM of ≥3 in independent experiments. When primary HNECs were used, all data were repeated in ≥3 separate patients.

## Results

Pyocyanin and 1-hydroxyphenazine are produced from the same precursor, Phenazine-1-carboxylicacid. The ratio of pyocyanin to 1-hydroxyphenzine production dictated by the balance of PhzM and PhzS enzymatic activity (Figure 1A). Both pyocyanin and 1-hydroxyphenazine have been shown to be excreted in lab grown *P. aeruginosa* strains during stationary phase^**30**^. We first investigated the ability of these molecules to increase cytosolic Ca^2+^ (Ca^2+^_cyt_) signaling by imaging submerged primary human nasal epithelial cells (submerged HNECs) and RPMI 2650 nasal squamous carcinoma cells loaded with Ca^2+^ indicator dye Fluo-4. Interestingly, pyocyanin induced slow-onset but sustained Ca^2+^_cyt_ responses in both cell types, while 1-hydroxyphenazine did not (Figures 1B-E). Mitochondria are important Ca^2+^ buffering organelles that can take up Ca^2+^ when Ca^2+^_cyt_ is elevated^**17**^. Similarly, pyocyanin, but not 1-hydroxyphenazine induced elevations of mitochondrial Ca^2+^ (Ca^2+^_mito_) in both RPMI 2650 and submerged HNECs (Figures 1F-I). However, the Ca^2+^_mito_ increase with pyocyanin was not significantly decreased by inhibitors of the outer membrane Ca^2+^ channels (voltage-dependent anion channel [VDAC1] inhibitor VBIT-12^**31**^) or inner membrane Ca^2+^ channels (mitochondrial Ca2+ uniporter [MCU] inhibitors benzethonium chloride^**32**^ or DS15670511^**33**^) in RPMI 2650s (Figures 1F-G). This is in contrast with our observations of 3-oxo-C12HSL-induced Ca^2+^_mito_ signaling, which is blocked by both benzethonium chloride and DS15670511. These data suggest that pyocyanin, unlike 3-oxo-C12HSL, increases mitochondrial Ca^2+^ through a non-MCU pathway.

**Figure 1.**
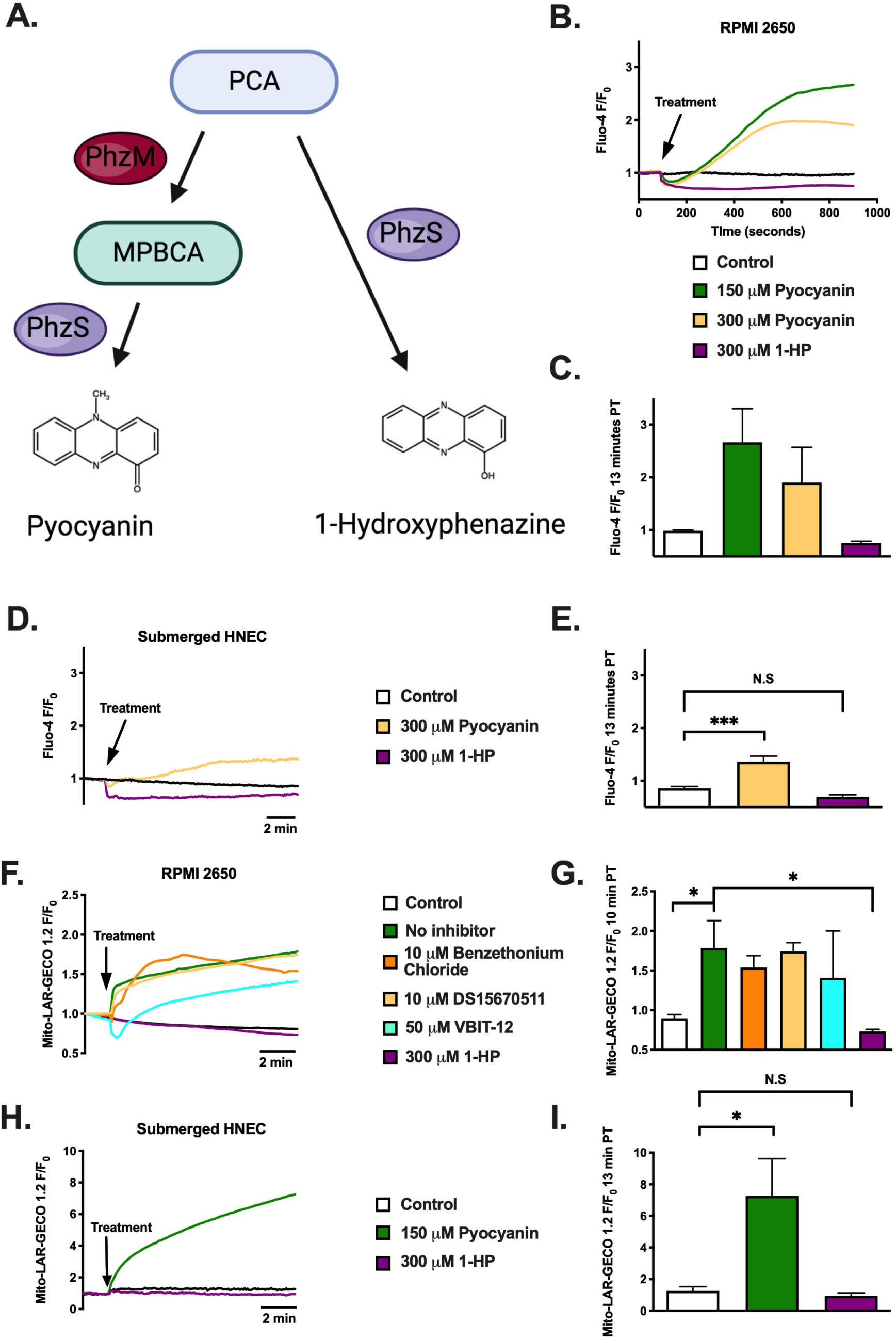
Pyocyanin induces cytoplasmic and mitochondrial Ca^2+^ responses in RPMI 2650 and submerged HNEC while 1-hydroxyphenazine does not. **A).** Phenazine-1-carboxylicacid (PCA) precursor is converted to 5-methylphenazine-1-carboxylic acid betaine (MPBCA) by the methyltransferase PhzM^**49**^, which is converted to pyocyanin by hydroxylase PhzS^**50**^. PhzS, in the absence of PhzM, or build up in PCA, directly converts PCA to 1-hydroxyphenazine^**51**^. **B).** Average traces of Fluo-4 AM intracellular Ca^2+^ responses in RPMI 2650 nasal squamous carcinoma cells. Responses were induced by 150 μM, 300 μM pyocyanin (Pyo) but not 300 μM 1-hydroxyphenazine (1-HP), which are concentrations that naturally occur in *Pseudomonas* biofilms. **C).** Intracellular Ca^2+^ (mean ± SEM) 13 minutes post treatment in RPMI 2650. **D).** Average traces of Fluo-4 AM intracellular Ca^2+^ in submerged HNEC. Responses were induced by 300 μM Pyo but not 300 μM 1-HP. **E).** Intracellular Ca^2+^ (mean ± SEM) 13 minutes post treatment in submerged HNEC. **F).** Average traces of mitochondrial Ca^2+^ in RPMI 2650’s transfected with Mito-LAR-GECO 1.2. Responses were induced by 150 μM Pyocyanin (Pyo) but not 300 μM 1-hydroxyphenazine (1-HP). This was not significantly different after pre-treatment with mitochondrial Ca^2+^ uniporter inhibitors 10 μM benzethonium chloride (Pyo w/BenzCl), or 10 μM DS16570511 (Pyo w/DS16), or with voltage dependent anion channel (VDAC) inhibitor 50 μM VBIT-12 (Pyo w/VBIT-12) HBSS and 1-HP treated wells were not pre-treated with any inhibitor and serve as controls **G).** Mitochondrial Ca^2+^ (mean ± SEM) 10 minutes post treatment in RPMI 2650s. **H).** Average traces of mitochondrial Ca^2+^ in submerged HNEC transfected with Mito-LAR-GECO 1.2. **I).** Mitochondrial Ca^2+^ (mean ± SEM) 13 minutes post treatment in submerged HNEC.

Fitting with increased PLC phospholipase C (PLC) activation and Ca^2+^ signaling, only pyocyanin increased protein kinase C (PKC) activity in RPMI 2650s while 1-hydroxyphenazine did not (Figures 2A-B). The activation of Ca^2+^_cyt_ by pyocyanin in RPMI 2650s was completely diminished by phospholipase C (PLC) inhibition (Figures 2C-D). Additionally, only pyocyanin caused endoplasmic reticular (ER) Ca^2+^ release in RPMI 2650 after stimulation, as measured by a fluorescent, ER-targeted cameleon biosensor, D1ER. In contrast, 1-hydroxyphenazine appeared to increase ER Ca^2+^ uptake (Figures 2E-F), though the mechanism of this was not interrogated. Nonetheless, these data together all suggest possible activation of one or more GPCRs by pyocyanin that stimulates PLC activation, IP3 generation, and ER Ca^2+^ release.

**Figure 2.**
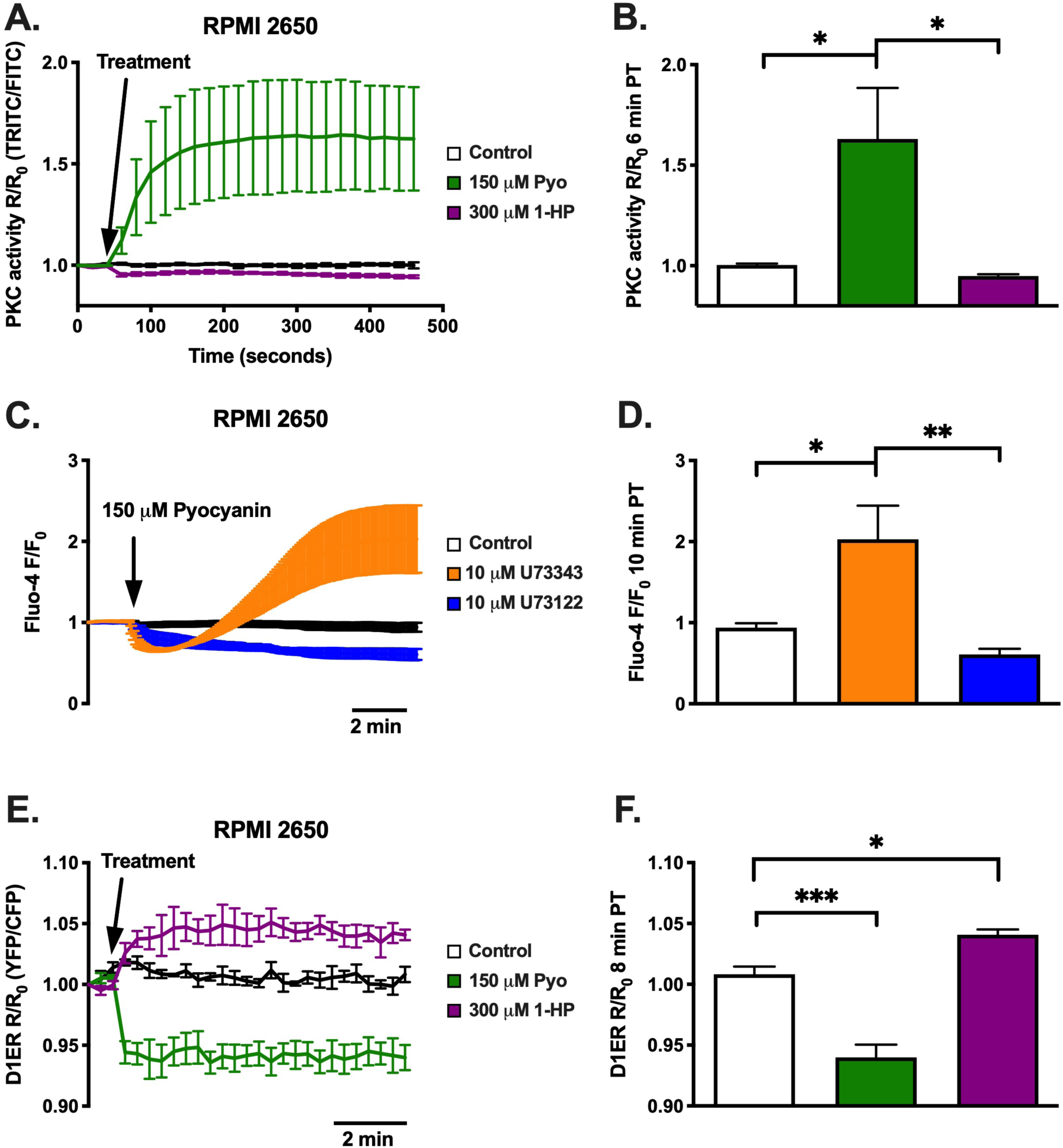
Pyocyanin induces PKC activation while 1-hydroxyphenazine does not and pyocyanin activation is diminished during PLC inhibition in RPMI 2650. **A).** Average traces, with error bars, of PKC activity in RPMI 2650s transfected with ExRai-CKAR. Responses were induced by 150 μM pyocyanin (Pyo) but not 300 μM 1-hydroxyphenazine (1-HP). **B).** PKC activity (mean ± SEM) 6 minutes post treatment in RPMI 2650s. **C).** Average traces, with error bars, of Fluo-4 intracellular Ca^2+^ responses in RPMI 2650 that were pre-treated 1 hour before with U73122 PLC inhibitor vs U73343 inactive control. 150 μM pyocyanin induced a calcium response in the inactive control, but no response in the active PLC inhibitor. **D).** Intracellular Ca^2+^ (mean ± SEM) 10 minutes post treatment in RPMI 2650. **E).** Average traces, with error bars, of ER Ca^2+^ activity in RPMI 2650s transfected with D1ER. Decreases in ER Ca^2+^ were seen with 150 μM pyocyanin, but was not seen with 300 μM 1-hydroxphenazine. **F).** ER Ca^2+^ activity (mean ± SEM) 8 minutes post treatment in RPM 2650s.

Despite the differential effects on Ca^2+^ signaling and PKC activation, mitochondrial membrane potential drastically decreased within minutes after treatment with either pyocyanin or 1-hydroxyphenazine. We tested this using both tetramethylrhodamine ethyl esterTMRE (single wavelength) or JC-1 (dual wavelength) mitochondrial membrane indicators in either submerged HNEC or RPMI 2650 (Figure 3). Pre-loading RPMI 2650 with potent Ca^2+^ chelator BAPTA (1-hour incubation with 10 µM BAPTA-AM) did not diminish mitochondrial membrane depolarization (Figure 3G-H), indicating that this depolarization is independent of Ca^2+^ signaling.

**Figure 3.**
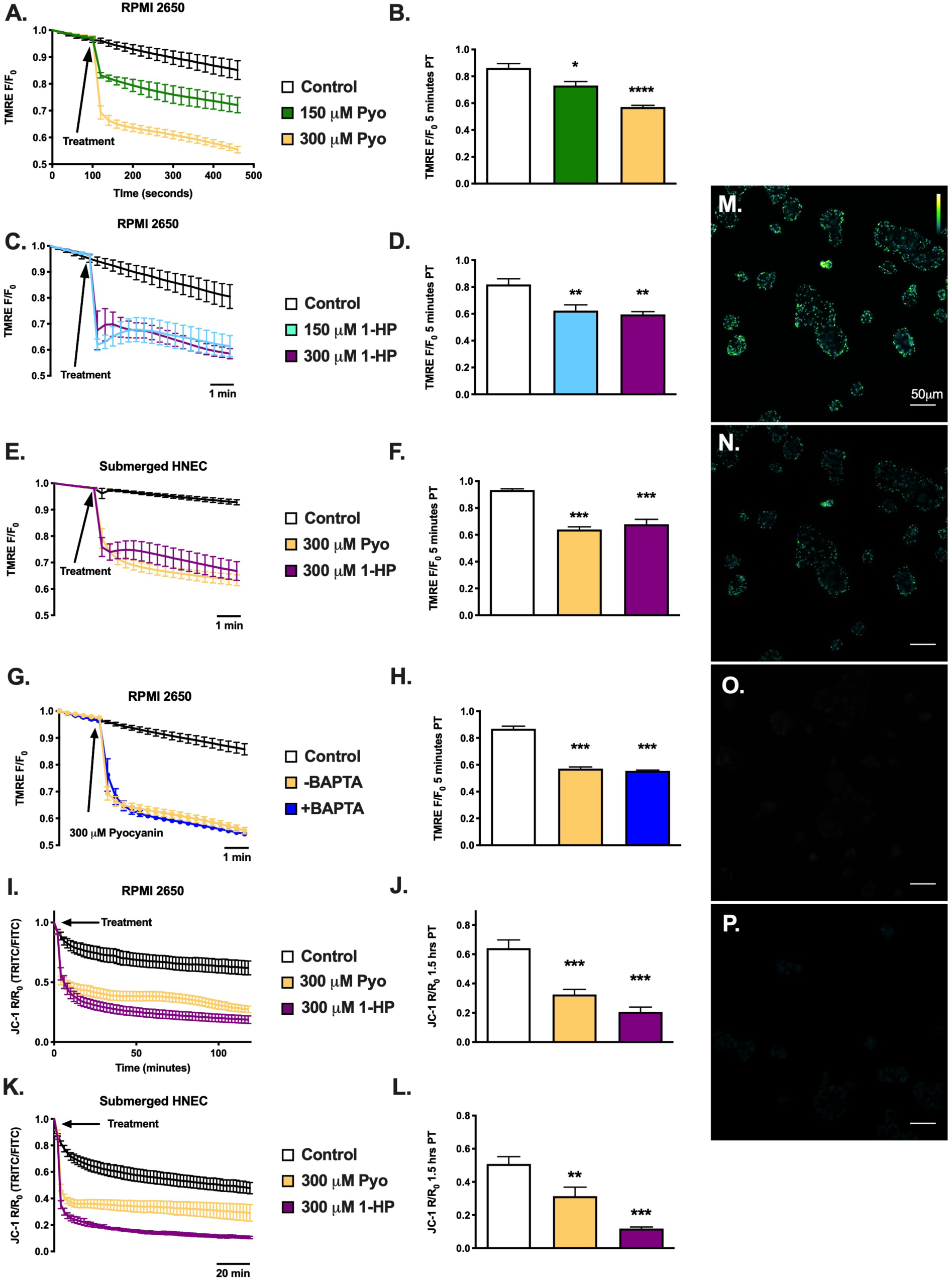
Acute mitochondrial depolarization in RPMI 2650 and submerged HNEC from pyocyanin and 1-hydroxyphenazine. **A).** Average traces, with error bars, of tetramethylrhodamine ethyl ester (TMRE) dye fluorescence over time and **B).** mean ± SEM after 4.5 minutes in RPMI 2650 after treatment with HBSS, 150 μM, and 300 μM pyocyanin. A reduction in TMRE fluorescence reflects a loss of mitochondrial membrane potential. **C).** Average traces, with error bars, of TMRE and **D).** mean ± SEM after 4.5 minutes in RPMI 2650 after treatment with HBSS (control), 150 μM, and 300 μM 1-hydroxyphenazine**. E).** Average traces, with error bars, of TMRE dye and **F).** mean ± SEM after 4.5 minutes in submerged primary cells (HNEC) after treatment with HBSS (control), 300 μM pyocyanin, and 300 μM 1-hydroxyphenazine **G).** Average traces, with error bars, of TMRE pretreated with or without (+/-) 10 μM BAPTA calcium chelator and **H).** mean ± SEM after 5 minutes in RPMI 2650 after treatment with HBSS (control), and 300 μM pyocyanin. **I).** Average traces, with error bars, of JC-1 dye fluorescence over time and **J).** mean ± SEM after 1.5 hours in RPMI 2650 after treatments with HBSS (control), 300 μM pyocyanin, and 300 μM 1-hydroxyphenazine. **K).** Average traces, with error bars, of JC-1 dye fluorescence over time and **L).** and mean ± SEM after 1.5 hours in Submerged HNEC after treatments with HBSS (control), 300 μM pyocyanin, and 300 μM 1-hydroxyphenazine. Similar to TMRE, the reduction in JC-1 R/R_0_ fluorescence indicates a loss of mitochondrial membrane potential. **M-P).** Representative images of JC-1 loaded RPMI 2650 (just TRITC) **M).** before treatment with HBSS (control), and 1.5 hours post treatment with **N).** HBSS, **O).** 300 μM pyocyanin, or **P).** 300 μM 1-hydroxyphenazine. Images were captured on a 20x objective and TRITC/FITC filters. All scale bars are 50 μm.

However, while pyocyanin, even in lower concentrations (10 μM), showed a strong increase in superoxide production in RPMI 2650 and seen especially in submerged HNECs (Figures 4A-D). Further pre-treatment with 10 μM BAPTA, similar to results from Figure 3, showed no significant decrease superoxide production in RPMI 2650 as a result of pyocyanin treatment after 2 hours, although BAPTA pre-treatment did slightly hinder 1-hydroxyphenazine superoxide production after 2 hours in the same cell line (Figures 4E-H). Visual representations show RPMI 2650 (Figures 4I-J) and submerged HNEC (Figures 4K-L) before (Figures I and K) and 2 hours after treatment with 100 μM pyocyanin (Figures J and L).

**Figure 4.**
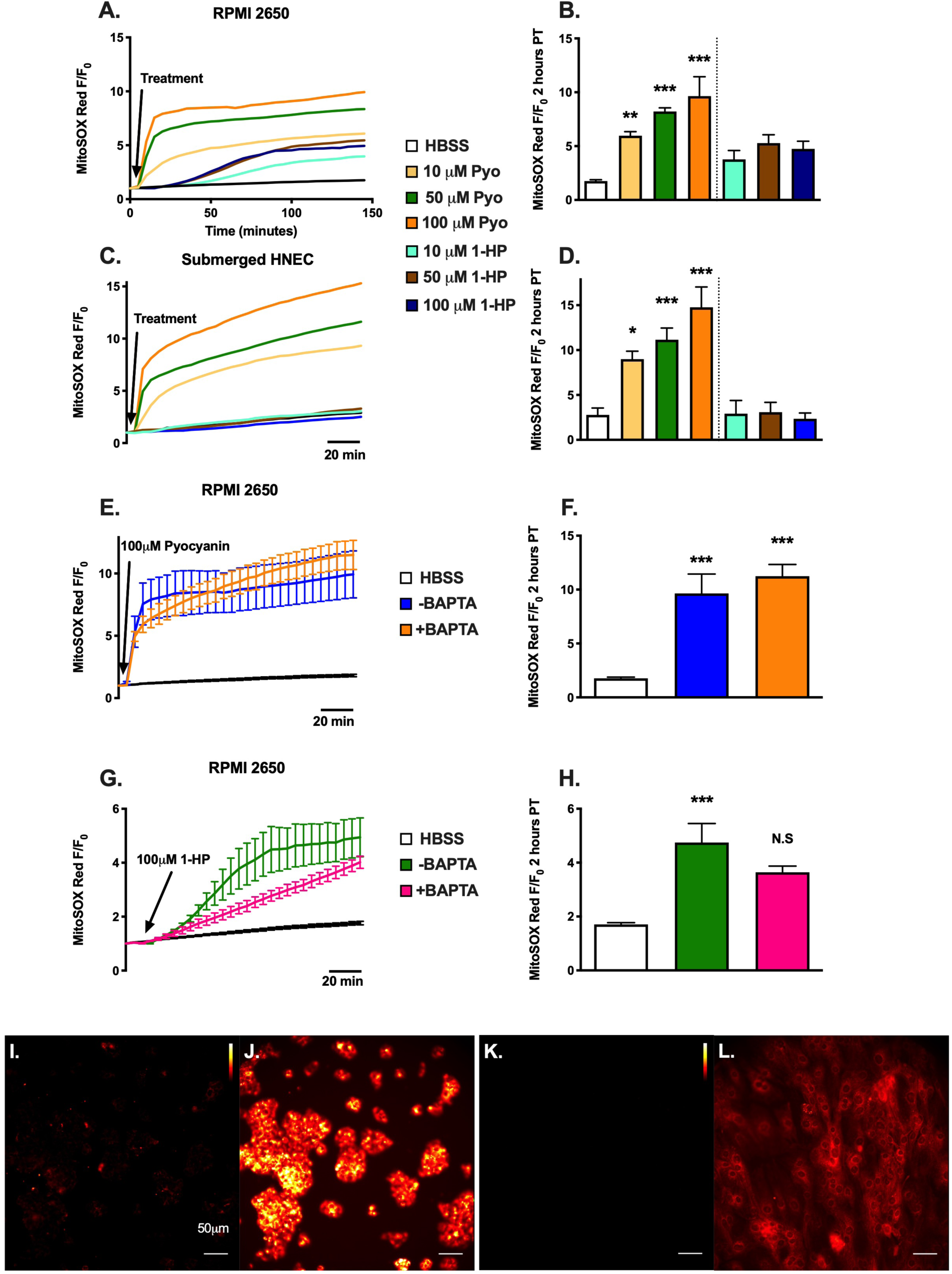
Pyocyanin and 1-hydroxyphenazine induce superoxide production in RPMI 2650 and but only pyocyanin induces superoxide production in primary nasal cells. **A).** Average traces of MitoSOX Red fluorescence over time and **B).** mean ± SEM after 2 hours in RPMI 2650 after treatment with HBSS (control), 100 μM pyocyanin, 50 μM pyocyanin, 10 μM pyocyanin, 100 μM 1-hydroxyphenazine, 50 μM 1-hydroxyphenazine, or 10 μM 1-hydroxyphenazine. An increase in MitoSOX Red fluorescence indicates an increase in superoxide production. **C).** Average traces of MitoSOX Red fluorescence over time and **D).** mean ± SEM after 2 hours in submerged primary nasal epithelial cells after treatment with HBSS, 100 μM pyocyanin, 50 μM pyocyanin, 10 μM pyocyanin, 100 μM 1-hydroxyphenazine, 50 μM 1-hydroxyphenazine, or 10 μM 1-hydroxyphenazine. **E and G).** Average traces, with error bars, of TMRE pretreated with or without (+/-) 10 μM BAPTA calcium chelator and **F and H).** mean ± SEM after 2 hours in RPMI 2650 after treatment with HBSS, **F).** 300 μM pyocyanin, or **H).** 300 μM 1-hydroxyphenazine. Representative images of MitoSOX Red loaded **I-J).** RPMI 2650 and **K-L).** submerged primary nasal epithelial cells **I and K).** at baseline, and **J and L).** 2 hours post treatment with 100 μM pyocyanin. Images were captured on a 20x objective and TRITC filter. All scale bars are 50 μm.

Nonetheless, both pyocyanin and 1-hydroxyphenazine decreased cell viability in squamous carcinoma cell lines RPMI 2650, SCC-47, and NCI-H520, but not in submerged HNECs (either from CF individuals or non-CF individuals) at 24 hours, shown by crystal violet staining (Figure 5). HNECs from three individual non-CF and three CF individuals appeared to be resistant to the cytotoxic effects of pyocyanin and 1-hydroxyphenazine.

**Figure 5.**
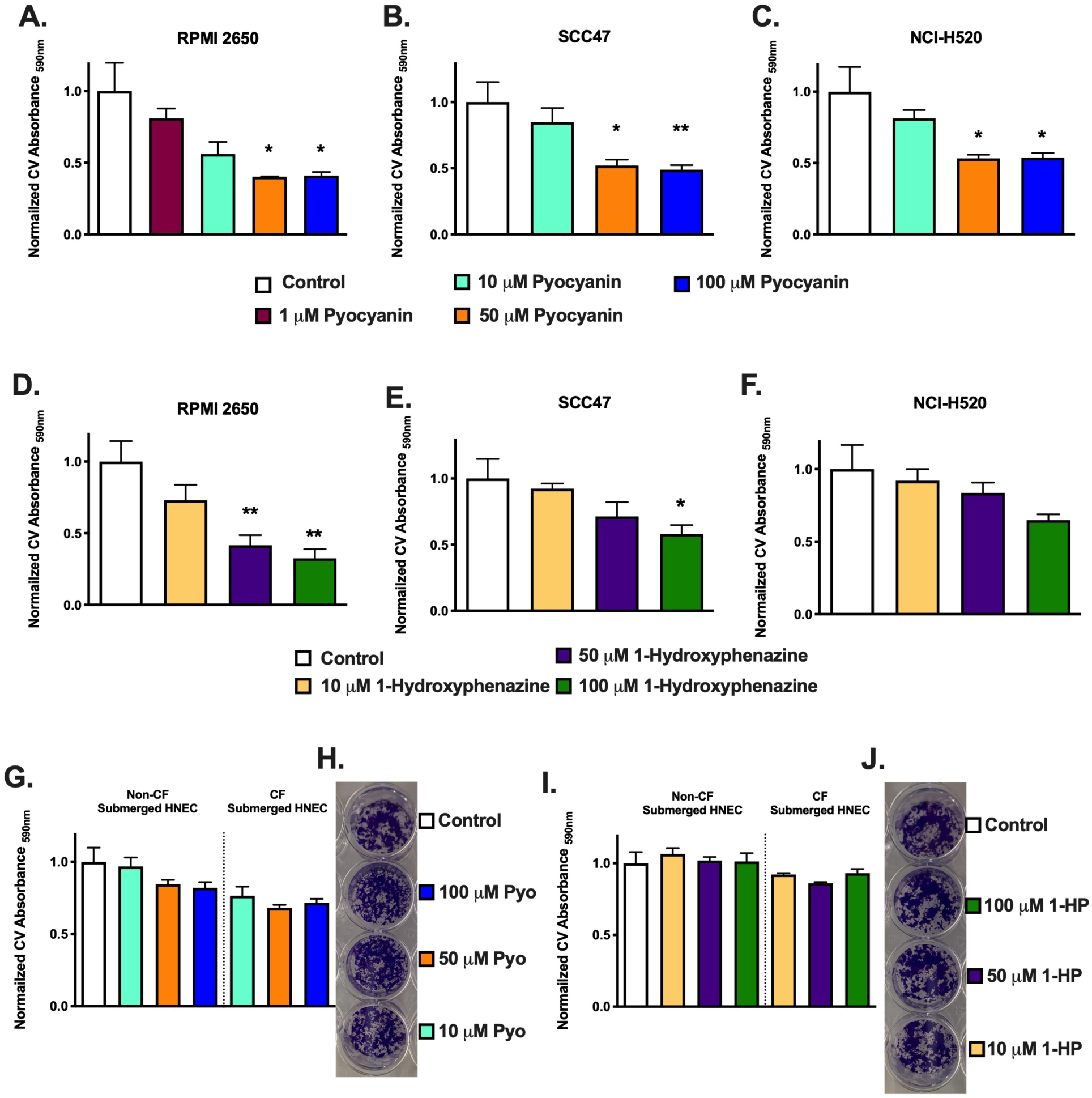
Pyocyanin and 1-hydroxyphenazine reduce viability of RPMI 2650 nasal carcinoma cells but not primary nasal epithelial cells. **A-C).** Crystal Violet (CV) absorbance in **A).** RPMI 2650s, **B).** SCC-47, and **C).** NCI-H520 all after 24-hour treatment with media (control), 100 μM, 50 μM, 10 μM, or 1 μM pyocyanin (only RPMI 2650). A decrease in CV absorbance indicates a reduction in cell numbers and thus cell viability. RPMI 2650, SCC-47, and NCI-H520 are all squamous cell carcinomas originating from the nasal septum, lateral tongue, and lung mass respectively. **D-F).** Crystal Violet (CV) absorbance in **D).** RPMI 2650 **E).** SCC-47, and **F).** NCI-H520 all after 24-hour treatment with media (control), 100 μM, 50 μM, or 10 μM 1-hydroxyphenazine. **G-J).** CV absorbance in submerged HNEC (non-CF vs CF), after 24-hour treatment with media, 100 μM, 50 μM, or 10 μM **G).** pyocyanin with **H).** representative image, or **I).** 1-hydroxyphenazine with **J).** representative image.

Interestingly pyocyanin did not induce robust Ca^2+^_cyt_ responses in either non-CF or CF HNECs differentiated at air liquid interface (ALIs), differing from what was seen in submerged HNECs and RPMI 2650s (Figure 6A-C). However, Ca^2+^_mito_ did increase in HNEC ALIs with pyocyanin, as measured via Rhod-2. Ca^2+^_mito_ also decreased when exposed to 1-hydroxyphenazine, similar to the effects seen in both submerged HNEC and RPMI 2650 (Figure 6D-E). It may be that the organization of ER/mitochondrial contact points in polarized ALI cultures allows more direct transfer of Ca^2+^ from the ER to the mitochondria, resulting in a reduced Ca^2+^_cyt_ responses but intact Ca^2+^_mito_ responses. Nonetheless, pyocyanin and 1-hydroxyphenazine both induced mitochondrial membrane depolarization, although the depolarization seen with pyocyanin was much greater than that of 1-hydroxyphenazine (Figure 6F-G).

**Figure 6.**
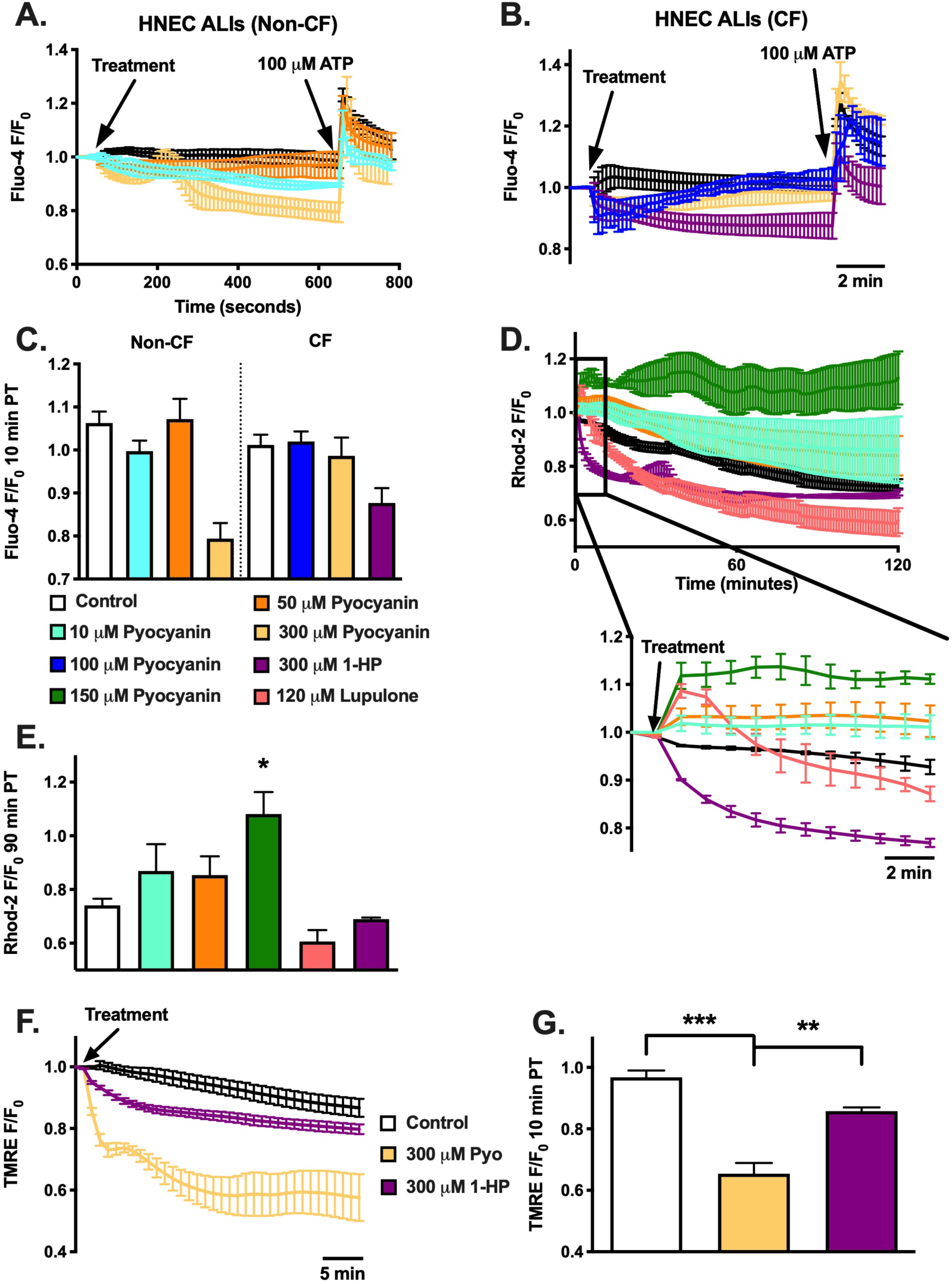
Pyocyanin and 1-hydroxyphenazine effects on primary HNEC ALI cytosolic and mitochondrial calcium, and mitochondrial membrane potential. **A-B).** Average traces, with error bars, of Fluo-4 AM intracellular Ca^2+^ in HNEC ALI **A).** (non-CF), **B).** (CF), and **C).** Intracellular Ca^2+^ (mean ± SEM) 10 minutes post treatment in HNEC ALI (non-CF vs CF) with HBSS (control), 10 μM, 50 μM, 100 μM, 300 μM pyocyanin, or 300 μM 1-hydroxyphenazine. No treatment was statistically significant at 10 minutes post treatment compared to any other treatment type **D).** Average traces, with error bars, of Rhod-2 AM mitochondrial Ca^2+^ in HNEC ALI (non-CF), and **E).** Intracellular Ca^2+^ (mean ± SEM) 90 minutes post treatment in HNEC ALI with HBSS (control), 10 μM, 50 μM, 100 μM pyocyanin, 120 μM lupulone, or 300 μM 1-hydroxyphenazine. Both 150 μM pyocyanin (***) and 120 μM lupulone (**) were significant, compared to HBSS, at 1-minute post treatment. **F).** Average traces, with error bars, of TMRE and **G).** mean ± SEM after 10 minutes in HNEC ALI (non-CF) after treatment with HBSS (control), 300 μM pyocyanin, or 300 μM 1-hydroxyphenazine. 300 μM pyocyanin was statistically significant at 10 minutes post treatment compared to both control (HBSS) and 300 μM 1-hydroxyphenazine.

Ciliary beat frequency (CBF) was decreased after 4 hours of exposure to 150 μM pyocyanin in HNEC ALIs, while 1-hydroxyphenazine had no effect (Figure 7A-B). Interestingly, pyocyanin induced superoxide production in both CF and non-CF HNEC ALIs, but CF cells used had significantly higher production compared to the non-CF cells (Figure 7C-D).

**Figure 7.**
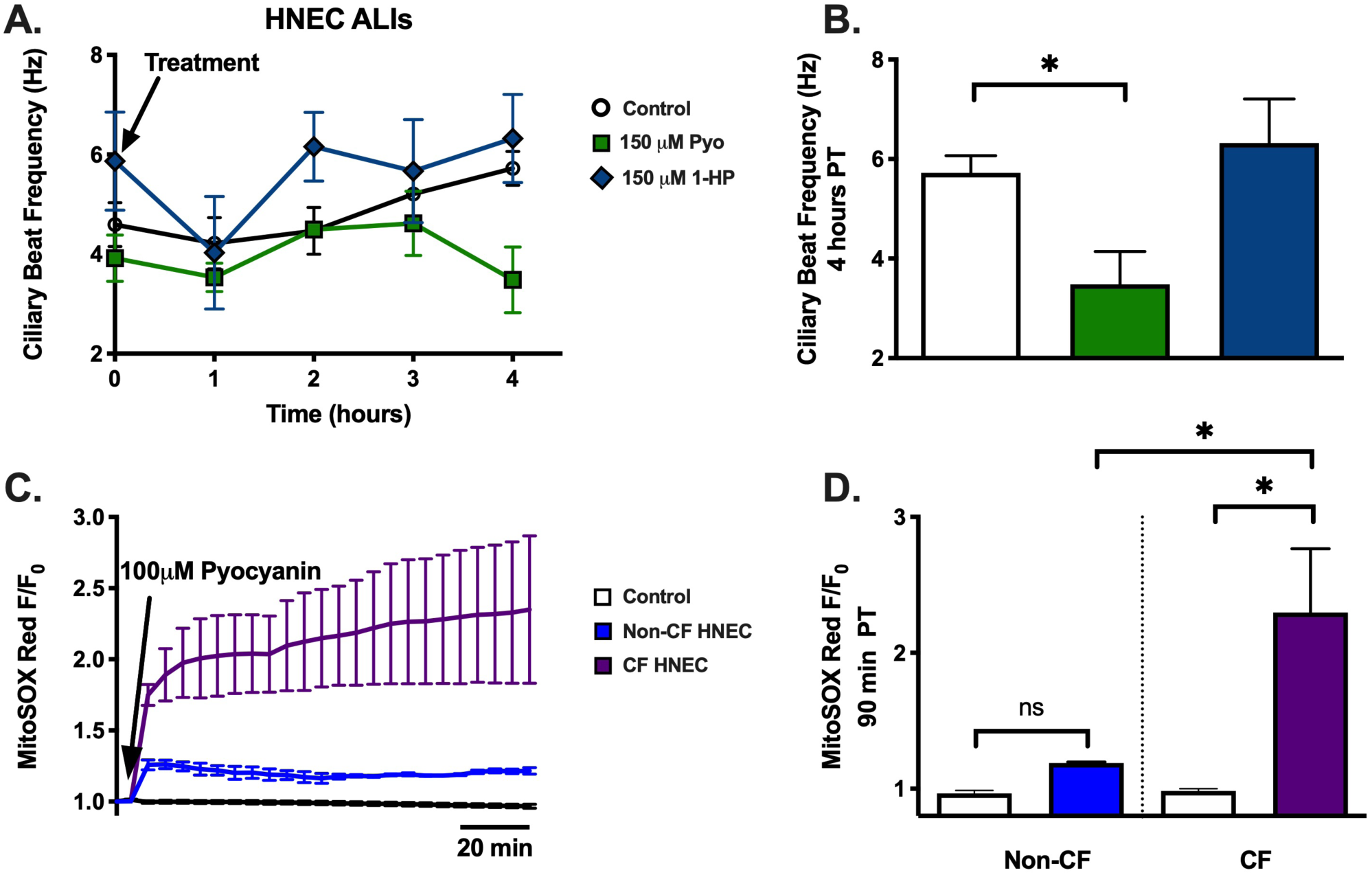
Pyocyanin induces decreases in ciliary beat frequency (CBF) and induce superoxide production in HNEC ALIs. **A).** Average traces, with error bars, of CBF (Hz) in HNEC ALI (non-CF) and **B).** CBF (Hz) (mean ± SEM) 4 hours post treatment in HNEC ALI (non-CF) after treatment with HBSS (control), 150 μM pyocyanin, or 150 μM 1-hydroxyphenazine. Only 150 μM pyocyanin was statistically significant at 4 hours post treatment when compared to control by two-tailed, unpaired t test. **C).** Average traces, with error bars, of MitoSOX Red fluorescence over time and **D).** mean ± SEM after 90 minutes in HNEC ALI (non-CF vs CF) after treatment with HBSS (control) or 100 μM pyocyanin. Only 150 μM pyocyanin (CF) was statistically significant at 90 minutes post treatment when compared to control or 150 μM pyocyanin (non-CF).

We hypothesized that pyocyanin depletion of ER Ca^2+^ might up-regulate the ER unfolded protein response (UPR^ER^). We looked at expression changes of gene involved in UPR^ER^ and ER-associated degradation (ERAD). Surprisingly, we found significant downregulation of ATF6, EDEM1, EDEM2, and EDEM3 in RPMI 2650s and SCC-47s but not in NCI-H520s and HNEC ALIs (Figure 8). The downregulation of genes involved ER protein homeostasis might be involved in some of the cytotoxic effects seen in RPMI 2650s and SCC-47s (Figure 5).

**Figure 8.**
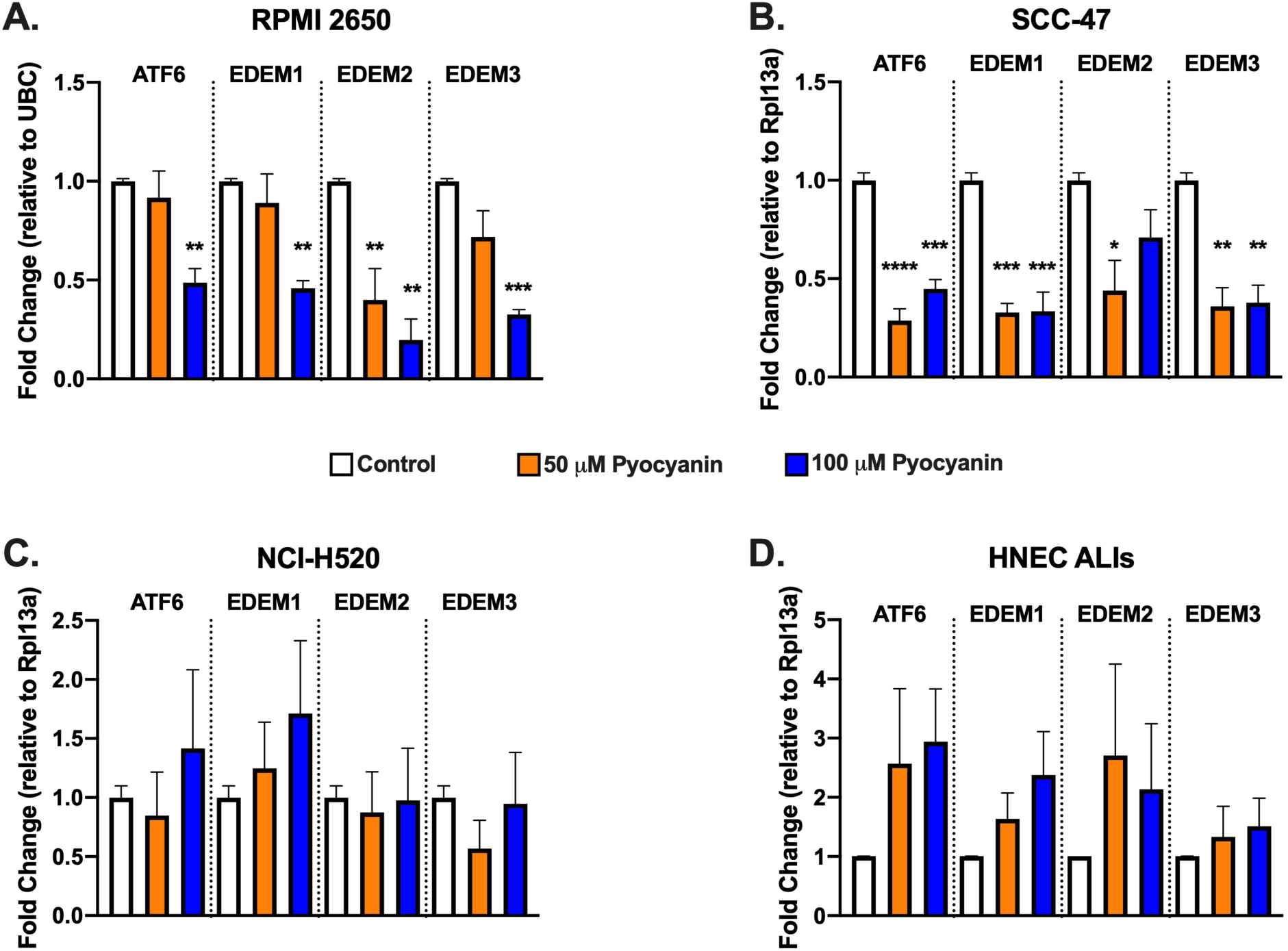
Pyocyanin induces ER stress response in some squamous cell carcinomas, but not primary HNEC. **A-D).** *ATF6, EDEM1, EDEM2,* and *EDEM3* expression in **A).** RPMI 2650, SCC-47, **C).** NCI-H520, or **D).** HNEC ALIs. Statistically significant decreases in all genes shown were seen in RPMI 2650 and SCC-47 with 100 μM pyocyanin and in SCC-47s with 50 μM pyocyanin when done by one-way ANOVA with Dunnett’s post-test. In either NCI-H520 or HNEC ALIs, gene expression was not statistically different.

## Discussion

P. aeruginosa plays a key role in lower airway infections in CF lung disease^**34**^ and upper airway infections in CF-related chronic rhinosinusitis^**35**^. Pyocyanin and 1-hydroxyphenazine are *P. aeruginosa* metabolites produced during P. aeruginosa infection. A myriad of effects have been reported for pyocyanin in airway epithelial cells, including reduced ciliary beating^**11,36**^, goblet cell metaplasia^**37**^ and IL-8 production^**38**^. Pyocyanin has been proposed to have toxic effects on mitochondrial by redox cycling with NADPH, generating superoxide (O(2)*), hydrogen peroxide (H(2)O(2)), and hydroxyl radical (HO*), which may directly oxidize glutathione^**39**^ and enhances NFκB activation during co-stimulation with TLR5 agonist flagellin^**40**^. Oxidative stress production by pyocyanin has been proposed to inhibit Cl-secretion via the cystic fibrosis transmembrane conductance regulator (CFTR)^**41**^ and inhibit the dual oxidase (DUOX)-thiocyanate (SCN^-^)-lactoperoxidase (LPO) antimicrobial defense system^**42**^. Pyocyanin-induced oxidative stress has also been proposed to underly the goblet cell metaplasia and mucus hypersecretion^**43**^.

Better understanding the effects of pyocyanin and 1-hydroxyphenazine on nasal epithelial cell mitochondrial function is important for understanding host-pathogen interactions during P. aeruginosa infection. We started investigating Ca^2+^ signaling, as pyocyanin has been linked to Ca^2+^ elevations^**18**^ ^**13**^. While Ca^2+^ is essential for oxidative phosphorylation and other mitochondrial metabolic pathways, excessive Ca^2+^ influx into the mitochondria in nasal epithelial cells can cause mitochondrial Ca^2+^ overload and excess ROS production^**7,23**^. We hypothesized that Ca^2+^ would be involved in mitochondrial toxicity observed with both pyocyanin and 1-hydroxyphenazine.

Here, we found that only pyocyanin induces Ca^2+^ responses, both cytosolic and mitochondrial in submerged HNEC and cancer cells, although no cytosolic Ca^2+^ response was seen in HNEC ALI, although they did show a prolonged mitochondrial response. 1-hydroxyphenazine had no detectible effects on Ca^2+^ in any of these cell types. Pyocyanin increased PKC activity, diminished cytosolic Ca^2+^ with a PLC inhibitor, and decreased ER Ca^2+^ stores indicating the potential activation of a GPCR. Despite the cytosolic and mitochondrial Ca^2+^ data, both pyocyanin and 1-hydroxyphenazine displayed decreases in cell viability in RPMI 2650s but not in submerged HNEC at 24 hours. However, in both RPMI 2650 and HNEC (submerged or ALI), mitochondrial membrane potential drastically decreased after treatment with either pyocyanin or 1-hydroxyphenazine. Furthermore, only pyocyanin greatly increased superoxide production in both RPMI 2650 and submerged HNEC, while 1-hydroxyphenazine had small increases or no effect. In HNEC ALI, CF cells showed a greater sensitivity to pyocyanin in superoxide production while non-CF cells had only small increases in superoxide production. HNSCC Our data suggest that the decrease in mitochondrial function and cell viability induced by both pyocyanin and 1-hydroxyphenazine is thus not likely through a T2R/Ca^2+^-dependent pathway. Furthermore, CF cells may have a higher sensitivity towards pyocyanin and be more negatively impacted in comparison to non-CF cells, given CF submerged HNEC had lowered cell viability compared to non-CF cells, and CF ALIs increased ROS production significantly compared to non-CF ALIs. Lastly, pyocyanin seems to reduce UPR^ER^ and ERAD markers in head and neck squamous cell carcinomas, but not in lung carcinoma or non-cancer primary cells. This may indicate a reduction/inhibition of these protein degradation responses in some cancer cells, but not in primary cells which may explain pyocyanin’s mechanism of action as a cancer therapeutic.

Other unknown targets of pyocyanin, and likely 1-hydroxyphenazine, exist in airway epithelial cells. In bronchial epithelial cells, pyocyanin induces IL-8 synthesis partially via TLR signaling and the aryl hydrocarbon receptor but also through unknown pathways^**44**^. Pyocyanin was shown to activate the EGFR-PI3K-AKT/MEK1/2-ERK1/2 MAPK pathway in A549 cells (alveolar type-II epithelial cells) to increase nuclear Nrf-2^**45**^, though these pathways are typically pro-survival pathways rather than cell death pathways. Notably, an early study reported that while 1-hydroxyphenazine and pyocyanin both inhibited mitochondrial respiration in intact cells, only 1-hydroxyphenzine inhibited respiration of isolated liver mitochondria^**46**^. These authors hypothesized that inhibition of mitochondrial function by pyocyanin may require intracellular demethylase conversion of pyocyanin to 1-hydroxyphenazine. Early studies have also reported that 1-hydroxyphenazine can interact with ubiquinone-cytochrome b within the mitochondrial electron transport chain^**47,48**^. It may indeed be that the Ca^2+^ responses observed here are due to pyocyanin activation of an unknown target, possibly a GPCR, while the mitochondrial depolarization and reduced viability of cancer cells we observe here requires intracellular conversion of pyocyanin to 1-hydroxyphenazine.

The greater sensitivity of RPMI 2650 cells vs primary cells suggests these compounds might have some applicability in treating nasal squamous carcinoma or other types of head and neck squamous carcinomas (HNSCCs). The further differences seen in UPR^ER^ and ERAD markers between HNSCCs (RPMI 2650 and SCC-47) and lung carcinoma (NCI-H520) show further differences between these cells likely derived from cancer vs normal tissue origins and support pyocyanin’s potential use as a cancer cell-specific cancer therapeutic.

Taken all together, these data suggest that acute Ca^2+^ signaling is not required for reductions in mitochondrial function, as both Ca^2+^-activating pyocyanin and Ca^2+^-independent 1-hydroxyphenazine decrease mitochondrial function. Likewise, the acute reductions in mitochondrial function coupled with superoxide production may not be the only mechanism for the decreased cell viability in cells exposed to *P. aeruginosa* phenazines, as similar depolarization was observed in both HNECs and squamous cell carcinomas but only the cancer cells died. The decrease in UPR^ER^ and ERAD markers pose a different mechanism for the decreased viability seen in the nasal and tongue carcinomas (RPMI 2650 and SCC-47 respectively) and lack thereof in primary cells (submerged HNEC), but still do not explain the decreased cell viability seen in the lung carcinoma (NCI-H520). The exact mechanism(s) of cell death with pyocyanin and 1-hydroxyphenazine remain to be determined, but data related to ER stress (UPR^ER^ and ERAD) could play a role in some cancer cells.

## Acknowledgements

This study was supported by National Institutes of Health grants HL168060 and AI167971 to R.J.L. The funders had no role in study design, conclusions, or decision to submit the manuscript. We thank M. Victoria (University of Pennsylvania) for excellent technical assistance.

